# Selection and evaluation of qPCR reference genes for expression analysis in the tiny egg parasitoid wasp, *Trichogramma dendrolimi* Matsumura (Hymenoptera: Trichogrammatidae)

**DOI:** 10.1101/2021.07.27.454008

**Authors:** Liang-xiao Huo, Xue-ping Bai, Wu-nan Che, Su-fang Ning, Lin Lv, Li-sheng Zhang, Jin-cheng Zhou, Hui Dong

**Affiliations:** College of Plant Protection, Shenyang Agricultural University, Shengyang, Liaoning, PR China; Institute of Plant Protection, Chinese Academy of Agricultural Sciences, Beijing, PR China

**Keywords:** qPCR, reference gene, biological control, egg parasitoid, *Trichogramma dendrolimi*

## Abstract

The egg parasitoid *Trichogramma* spp. is an important biological control agent used against multiple species of Lepidopteran pest in forestry and agriculture. Due to the importance of *Trichogramma* spp. in biocontrol programs, its biological characteristics have been studied in detail, and current investigations should focus on the molecular biology of these tiny parasitoids. Real-time quantitative PCR (qPCR) is considered as the standard method for quantifying the gene expression of organisms. Surprisingly, the appropriate reference genes to ensure robust qPCR have not been documented at all for the *Trichogramma* genus. This study aimed to identify suitable reference genes for use in qPCR procedure of *Trichogramma dendrolimi*. Nine candidate housekeeping genes, namely glyceraldehyde-3-phosphate dehydrogenase (*GAPDH*), forkhead box O (*FOXO*), superoxide dismutase (*SOD*), beta-actin (*ACTIN*), ribosomal protein L10a (*RPL10a*), L18 (*RPL18*), L28 (*RPL28*), S13 (*RPS13*), and S15 (*RPS15*), were tested for their suitability as reference genes for developmental stage (3^rd^, 4^th^, 5^th^, 6^th^, 7^th^, 8^th^, 9^th^, and 10^th^ day after parasitization), tissue (head, thorax, and abdomen of adults), sex of adults (male and female), and temperature (17 °C, 25 °C, and 32 °C). According to the GeNorm analysis, robust analysis should involve using an appropriate combination of reference genes, namely, at least three genes for different development stages, two genes for different tissues, two genes for different sex, and two genes for different temperature, respectively. According to the RelFinder method and by assessing the integrated values from using the ΔCt method, GeNorm, NormFinder, and BestKeeper, we identified the developmental stage-specific reference genes *SOD, GAPDH*, and *ACTIN*; tissue-specific reference genes *RPL18* and *RPS15*; sex-specific reference genes *SOD* and *RPL18*; and temperature-specific reference genes *RPL18* and *RPL10*. When testing the use of stable vs. unstable reference genes, the substantial differences were observed in the estimation expression of a hypothetical target gene, *HSP90*, in response to temperature. The present study provides a robust method for the measurement of gene expression in *T. dendrolimi* and will be helpful for future biological control programs using *Trichogramma* wasps.

## Introduction

The egg parasitoid *Trichogramma* spp. has been viewed as an effective biological control agent against multiple species of Lepidopteran pests in forestry and agriculture [1,2]. Among all the *Trichogramma* species, *Trichogramma dendrolimi* is considered an important egg parasitoid of a broad range of pests [3–6]. In China, the use of *T. dendrolimi* for controlling Lepidopteran pests is particularly inexpensive due to it being reared on the eggs of Chinese oak silk moth (*Antheraea pernyi*), which is easy to obtain from farmers who rear and collect cocoons of *A. pernyi* for the textile industry and can support the development of approximately in the tens to hundreds of offspring wasps per egg [7–10]. Therefore, the inundative release of *T. dendrolimi* wasps is an increasingly important, efficient, and environmentally friendly measure to control Lepidopteran pests that has been applied across millions of hectares annually in the crop-producing areas of China [2].

As important biological control agents, the biological characteristics of *Trichogramma* spp. have been studied in detail [2,10,11], and current investigations should focus on the molecular biology of *Trichogramma*. Although the whole genomes of *Trichogramma*, including *T. pretiosum, T. brassicae*, and *T. evanescens*, have been well annotated [12–14], the regulatory mechanisms of these tiny parasitoids is poorly researched.

Real-time quantitative PCR (qPCR) has been viewed as the model method for quantifying gene expression of organisms [15,16]. Surprisingly, appropriate reference genes for the estimation and validation of gene expression have not been documented at all for the *Trichogramma* genus. qPCR results are easily influenced by multiple factors, especially in the samples of *Trichogramma* spp. [17,18]. First, the body sizes of *Trichogramma* wasps and immature offspring are extremely small (length of wasp is approximately 0.5 mm) [1]. These tiny parasitoids are highly sensitive to biotic or abiotic factors which result in fluctuations in gene expression. Due to the very low amounts of mRNA of the target gene in the tissues, their quality can also be easily affected throughout the whole RNA extraction procedure. Second, egg parasitoids, such as *Trichogramma* spp., develop internally in host eggs. The RNA quality of samples is often influenced by other residual content from host insects [19,20]. Additionally, the RNA from *Trichogramma* offspring may degrade during the isolation of the parasitoid body from host eggs. To eliminate errors in expression quantification resulting from the discussed reasons, it is critical to select suitable genes for normalizing qPCR results and establishing a robust method for quantifying the expression of target genes [21].

The appropriate reference genes should have ubiquitous expression with low variation and reasonable stability under different conditions [22]. Housekeeping genes are often applied as reference genes due to their stable and constant expression in all cells of the organism [21]. The mostly widely used genes for normalizing the expression of target genes include beta-actin (*ACTIN*) [23], glyceraldehyde-3-phosphate dehydrogenase (*GAPDH*) [24], ribosomal protein L10 (*RPL10*), L18 (*RPL18*), L28 (*RPL28*), S13 (*RPS13*), S15 (*RPS15*) [25], and superoxide dismutase (*SOD*) [26], because the constant expression of these genes are essential for the survival of organisms and they are synthesized in all types of cells [27]. Therefore, these genes have been used as reference genes in qPCR procedures of various organisms [28–30]. In hymenopteran insects, the identification of reliable reference genes has been conducted in the parasitoid wasps *Cotesia chilonis* [30], *Aphidius gifuensis* [19], and *Lysiphlebia japonica* [20], and in the bee species *Bombus terrestris* [31], *B. lucorum* [32], *Frieseomelitta varia* [33], *Melipona quadrifasciata* [33], *Scaptotrigona bipunctata* [33], *Apis mellifera* [34], and *Megachile rotundata* [35]. Given the importance of *T. dendrolimi* in biological control programs, the identification of suitable reference genes is considered necessary for extensive study on the molecular biology of *T. dendrolimi* and other *Trichogramma* species.

The present study was aimed to select suitable reference genes for use in qPCR procedure of *T. dendrolimi*. Nine housekeeping genes (*ACTIN*, forkhead box O (*FOXO*) [36,37], *GAPDH, RPL10a, RPL18, RPL28, RPS13, RPS15*, and *SOD*) were used as candidate reference genes and the potential influence on their expression according to different factors such as the developmental stage (3^rd^, 4^th^, 5^th^, 6^th^, 7^th^, 8^th^, 9^th^, and 10^th^ day after parasitization), tissue (head, thorax, and abdomen of adults), sex of adults (male and female), and temperature (25°C as the control, 17°C and 32°C) was evaluated. The stability of candidate genes expression was identified according to five statistical algorithms (the ΔCt method, GeNorm, NormFinder, BestKeeper, and RefFinder). The hypothetical target gene, heat shock protein 90 (*HSP90*), was applied to evaluate the accuracy of the reference genes. The results will facilitate improved qPCR analysis of this parasitoid species as influenced by biotic and abiotic factors.

## Materials and methods

### Insects

The isofemale line of *T. dendrolimi* was maintained in the biological control laboratory of College of Plant Protection in Shenyang Agricultural University. *T. dendrolimi* was reared on the eggs of *Antheraea pernyi* (Lepidoptera: Saturniidae). Eggs of *A. pernyi* were collected from the dissected ovaries of female moths that had emerged from cocoons. The extracted eggs were cleaned with ultrapure water, and nonviable eggs (green eggs and dead eggs) were abandoned. Eggs of *A. pernyi* (in groups of 50) were then glued on the strip-shaped cards (length × width: 80 mm × 5 mm) with white glue. The egg card was transferred into a plastic tube (120 mm length, 20 mm diameter, stoppered with cotton balls). A group of 200 *T. dendrolimi* wasps (1-day-old) was introduced into the glass and allowed to parasitize for 1 h, after which the wasps were removed. The parasitized eggs of *A. pernyi* were used in this experiment.

All insects were reared under 25 ± 0.5°C, 16 h/8 h (light/dark) photoperiod, and 70% ± 5% relative humidity.

### Sample preparation

Biotic and abiotic factors were considered in this study, including development stages (3^rd^, 4^th^, 5^th^, 6^th^, 7^th^, 8^th^, 9^th^, and 10^th^ day after parasitization), tissue (head, thorax, and abdomen), sex of adults (male and female), and temperature (25°C as the control and 17°C and 32°C).

The samples of *T. dendrolimi* offspring (in groups of 250 offspring individuals) were dissected and collected from parasitized eggs of *A. pernyi* at the 3rd (larvae), 4th (prepupae), 5th (prepupae), 6th (prepupae), 7th (pupae), 8th (pupae), 9th (pupae), and 10th (pupae) day after parasitization under a stereoscope (SMZ-161; Motic, Xiamen, China) [38]. The samples of head, thorax, and abdomen (in groups of 50) were dissected and isolated from the newly emerged females using a tungsten needle (tip diameter = 0.05 mm). The samples of females and males were determined based on recognition of the secondary sexual characteristics of *T. dendrolimi* and collected in groups of 30 individuals. To determine the effects of temperature, the newly emerged females, in groups of 30, were exposed to 17°C, 25°C, and 32°C for 2 h in Durham glass tubes (30 mm length, 6 mm diameter, stoppered with cotton balls).

All the samples were quickly transferred into 1.5 mL tubes and stored at −80°C. Each treatment or level involved three replicates.

### RNA extraction and cDNA synthesis

Total RNA of the samples was extracted by using TRIzol reagent (Ambion, Life Technologies, USA) according to the instructions of the manufacturer. The quality and concentration of RNA extractions were identified using a spectrophotometer (NanoDrop 2000, Thermo Fisher Scientific, USA) according to the value of A260/280, which ranged from 1.8 to 2.0. After adjusting the samples to have equal concentrations, 1.0 mg RNA was reverse-transcribed into first-strand cDNA using the PrimeScriptTMRT reagent Kit with the gDNA Eraser following the protocol of manufacturer (Takara, Dalian, China). The reverse transcriptional procedure included the following: 2 µL of 5× gDNA Eraser Buffer, 1 µg total RNA, 1 µL gDNA Eraser, RNase-free ddH_2_O up to 10 µL, with reaction at 42°C for 120 s, then adding 1 µL RT Primer Mix, 1 µL PrimerScript® RT-Enzyme Mix I, 4 µL of 5× PrimerScript^®^ Buffer 2, and RNase-free ddH_2_O up to 20 µL, with reaction at 37°C for 15 min, followed by reaction termination at 85°C for 5 s. The cDNA of all samples was stored at −20°C.

### Gene sequences and Primer design

Eight candidate reference genes (*ACTIN, GAPDH, SOD, RPL18, RPL28, RPL10a, RPS13*, and *RPS15*) and a hypothetical target gene (*HSP90*) were identified from the previously constructed RNA-seq transcriptome dataset of *T. dendrolimi*. The candidate reference gene, *FOXO*, was obtained from the GenBank database (accession number: KX066243). The ORF Finder was applied to identify the open reading frame (ORF) of the gene (http://www.ncbi.nlm.nih.gov/gorf/gorf.html). The primers of all genes which were used for the subsequent qPCR procedure were designed using NCBI Primer-BLAST (https://www.ncbi.nlm.nih.gov/tools/primer-blast/)(Table 1).

**Table 1.**
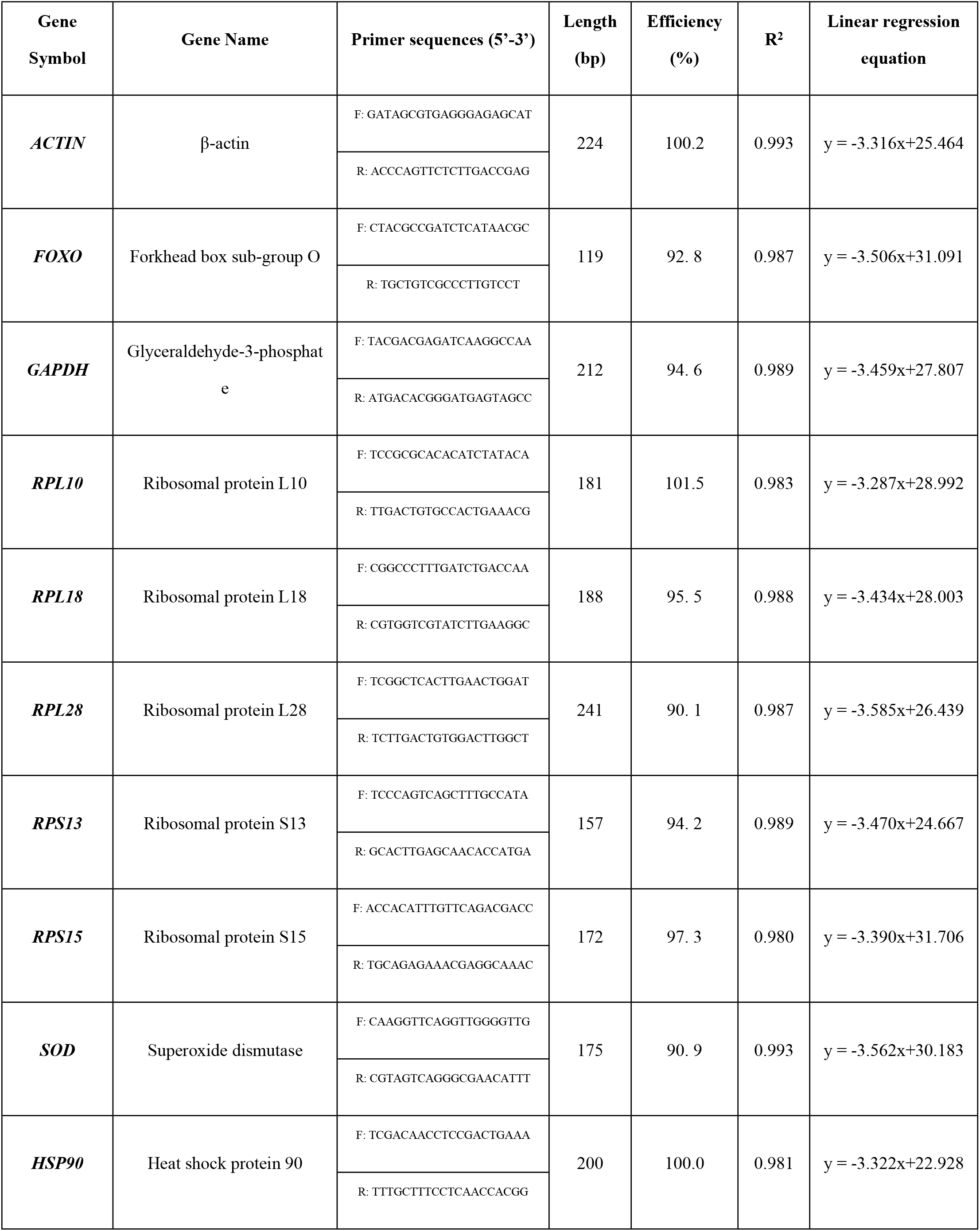
Sequence and amplification characteristics of qPCR primers for nine candidate reference genes and one hypothetical target gene.

### Procedure of RT-qPCR

The reactions of qPCR were conducted in the Bio-Rad CFX96 Real-time PCR Detection System (Bio-Rad, Hercules, California, USA) using 20 μL solutions containing 1.6 μL primers, 1.6 μL cDNA template, 10 μL of 2×SYBR Green Premix (Takara, Japan), and 6.8 μL ddH_2_O. The procedure of amplification was as follows: 95°C for 30 s, 40 cycles of 95°C for 5 s and 57°C for 30 s. Melting curve data were gathered from 57 to 95 °C. The standard curve was generated by a three-fold dilution series of cDNA [39]. Three technical replicates per sample were conducted independently.

### Data analysis of candidate reference genes

The constancy and stability of the genes was conducted independently for four factors: developmental stage, tissue, sexes of adults, and temperature, and with the following statistical algorithms: the ΔCt method, NormFinder, GeNorm, and BestKeeper.

GeNorm was applied to select the optimal reference gene by calculating a stability value (M) and determine the appropriate number of the combination of reference genes by comparing the pairwise variation (V).

The larger the M value is, the worse the stability and vice versa. The optimal number of the combination of reference genes was identified according to the V value of the certain factor after the introduction of a new gene, which was indicated by Vn/Vn+1. If Vn/Vn+1 is lower than 0.15, the appropriate number of genes should be “n”; otherwise, “n+1” is recommended [40].

NormFinder is used for ranking the constancy of candidate reference genes [17]. The calculation principle of NormFinder is similar to that of GeNorm. The optimal reference gene was also selected based on a stable parameter of the candidate genes. The criterion of NormFinder is the same as GeNorm.

BestKeeper is used for estimating the stability of candidate genes. The stability was finally determined by comparing the value of standard deviation (SD), correlation coefficient (r), and coefficient of variation (CV). The criterion is that the larger the r, the smaller the CV and SD and the better the constancy of the candidate gene. When SD > 1, the expression of the gene is unstable [41].

The ΔCt method was used to compare the stability of genes within the sample. The ranks are based on the value of ΔCt [42].

Finally, the constancy of candidate genes was ranked by RefFinder (http://blooge.cn/RefFinder/) according to the integrating results of the ΔCt method, GeNorm, NormFinder, and BestKeeper [43].

### Validation of selected reference gene

The hypothetical target gene, *HSP90*, was applied to verify the stability of the selected reference genes base on the ΔCt method, GeNorm, NormFinder, and BestKeeper. The expression of *HSP90* at 17°C, 25°C, and 32°C was calculated by the 2^−ΔΔCt^ method [44]. One-way ANOVA was used to analyze the expression of *HSP90* relative to different reference genes. The Tukey–Kramer test was used for multiple comparisons. The analysis was conducted using R 4.1.0 software [45].

## Results

### Primer Amplification efficiency

The specificity of primers was verified by 1.5% agarose gel electrophoresis. Standard curve analysis indicated that the efficiencies of primer amplification ranged from 90% to 105%, with high correlation coefficients (R^2^ > 0.98) (Table 1). In addition, The specificity of the primers were identified according to the presence of the single peak in the melting curve of each gene.

### Expression profiling of the candidate genes

The raw Ct values of nine candidate genes and one hypothetical target gene ranged from 21.16 (*HSP90*) to 31.48 (*FOXO*) in all samples, from 22.70 (*RPS13*) to 31.48 (*FOXO*) among the developmental stages, from 22.58 (*HSP90*) to 30.40 (*FOXO*) in females and males, from 21.97 (*HSP90*) to 29.85 (*FOXO*) among tissues, and from 21.16 (*HSP90*) to 29.28 (*FOXO*) at different temperatures, respectively (Fig. 1).

**Fig. 1.**
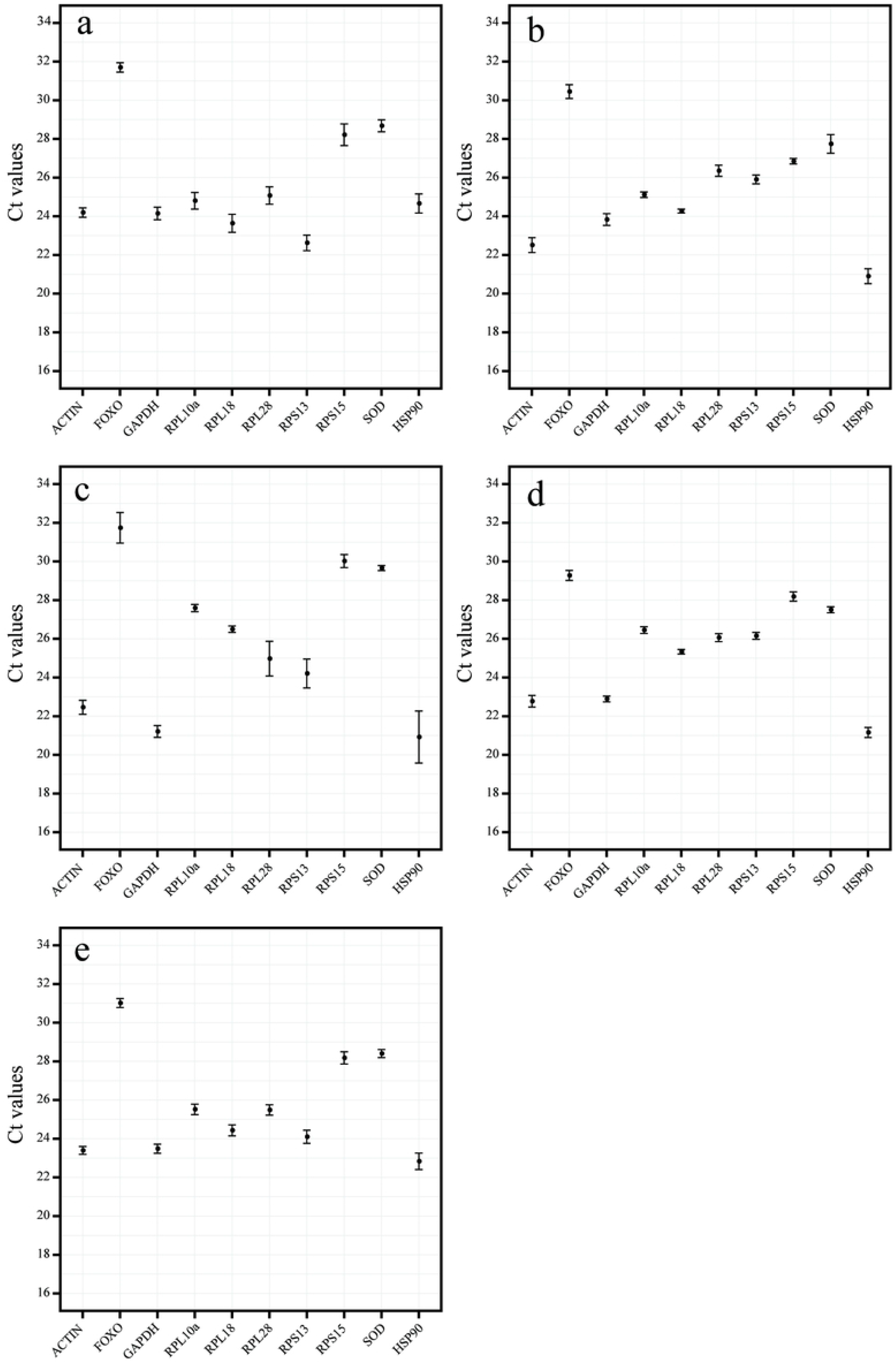
Expression profiles of the nine candidate reference genes and the hypothetical target gene *HSP90* in *T. dendrolimi* under different developmental stage (a), tissue (b), sexes of adult (c), and temperature (d). Expression levels are displayed as cycle threshold (Ct) values (mean ± SEM).

### Constancy of candidate genes

#### Developmental stage

According to results of GeNorm, the top three stable genes were *SOD, GAPDH*, and *RPL28*, and the top three unstable genes were *RPS13, FOXO*, and *RPS15*. According to the ΔCt method and NormFinder, the top three stable genes in terms of expression levels as influenced by developmental stages were *SOD, GAPDH*, and *ACTIN*, and the top three unstable genes were *RPS13, FOXO*, and *RPS15*. According to the analysis of BestKeeper, the top three stable genes were *FOXO, ACTIN*, and *SOD*, and the top three unstable genes were *RPS15, RPL18*, and *RPL28* (Table 2).

**Table 2.** Stability of candidate genes as influenced by different factor.

| Experimental conditions | Reference genes | GeNorm | | NormFinder | | BestKeeper | | $\Delta Ct$ | |
| --- | --- | --- | --- | --- | --- | --- | --- | --- | --- |
|  |  | Stability | Rank | Stability | Rank | Stability | Rank | Stability | Rank |
| Developmental stage | *ACTIN* | 1.248 | 6 | 0.862 | 3 | 0.857 | 2 | 1.493 | 3 |
|  | *FOXO* | 1.506 | 8 | 1.745 | 8 | 0.792 | 1 | 2.044 | 8 |
|  | *GAPDH* | 0.566 | 1 | 0.722 | 2 | 1.245 | 4 | 1.409 | 2 |
|  | *RPL10* | 1.175 | 4 | 0.882 | 4 | 1.513 | 6 | 1.512 | 4 |
|  | *RPL18* | 1.198 | 5 | 1.071 | 6 | 1.773 | 8 | 1.595 | 6 |
|  | *RPL28* | 1.023 | 3 | 0.941 | 5 | 1.579 | 7 | 1.521 | 5 |
|  | *RPS13* | 1.679 | 9 | 2.023 | 9 | 1.305 | 5 | 2.285 | 9 |
|  | *RPS15* | 1.342 | 7 | 1.591 | 7 | 1.998 | 9 | 1.900 | 7 |
|  | *SOD* | 0.566 | 1 | 0.585 | 1 | 1.154 | 3 | 1.355 | 1 |
| Tissue | *ACTIN* | 0.851 | 8 | 0.736 | 8 | 0.825 | 8 | 0.959 | 8 |
|  | *FOXO* | 0.790 | 7 | 0.682 | 6 | 0.801 | 7 | 0.941 | 6 |
|  | *GAPDH* | 0.639 | 5 | 0.710 | 7 | 0.655 | 6 | 0.945 | 7 |
|  | *RPL10* | 0.405 | 3 | 0.558 | 3 | 0.260 | 2 | 0.820 | 3 |
|  | *RPL18* | 0.356 | 1 | 0.412 | 1 | 0.203 | 1 | 0.761 | 1 |
|  | *RPL28* | 0.701 | 6 | 0.645 | 4 | 0.567 | 5 | 0.919 | 4 |
|  | *RPS13* | 0.553 | 4 | 0.677 | 5 | 0.424 | 4 | 0.925 | 5 |
|  | *RPS15* | 0.356 | 1 | 0.467 | 2 | 0.293 | 3 | 0.781 | 2 |
|  | *SOD* | 0.905 | 9 | 0.955 | 9 | 1.045 | 9 | 1.096 | 9 |
| Sex | *ACTIN* | 0.758 | 6 | 0.589 | 4 | 0.633 | 6 | 1.199 | 5 |
|  | *FOXO* | 1.368 | 9 | 1.890 | 9 | 1.539 | 9 | 1.965 | 9 |
|  | *GAPDH* | 0.648 | 5 | 0.601 | 5 | 0.417 | 4 | 1.183 | 4 |
|  | *RPL10* | 0.373 | 1 | 0.202 | 3 | 0.298 | 3 | 1.031 | 3 |
|  | *RPL18* | 0.373 | 1 | 0.131 | 2 | 0.251 | 2 | 1.030 | 2 |
|  | *RPL28* | 1.198 | 8 | 1.834 | 8 | 1.524 | 8 | 1.952 | 8 |
|  | *RPS13* | 1.010 | 7 | 1.520 | 7 | 1.440 | 7 | 1.713 | 7 |
|  | *RPS15* | 0.584 | 4 | 0.795 | 6 | 0.596 | 5 | 1.213 | 6 |
|  | *SOD* | 0.438 | 3 | 0.045 | 1 | 0.224 | 1 | 1.027 | 1 |
| **Temperature** | *ACTIN* | 0.669 | 9 | 0.712 | 9 | 0.405 | 6 | 0.835 | 9 |
|  | *FOXO* | 0.622 | 8 | 0.669 | 8 | 0.526 | 8 | 0.794 | 8 |
|  | *GAPDH* | 0.499 | 5 | 0.394 | 4 | 0.312 | 3 | 0.624 | 4 |
|  | *RPL10* | 0.383 | 1 | 0.275 | 2 | 0.354 | 5 | 0.566 | 2 |
|  | *RPL18* | 0.383 | 1 | 0.187 | 1 | 0.231 | 1 | 0.528 | 1 |
|  | *RPL28* | 0.471 | 4 | 0.400 | 5 | 0.584 | 9 | 0.628 | 5 |
|  | *RPS13* | 0.423 | 3 | 0.374 | 3 | 0.341 | 4 | 0.611 | 3 |
|  | *RPS15* | 0.578 | 7 | 0.617 | 7 | 0.478 | 7 | 0.760 | 7 |
|  | *SOD* | 0.532 | 6 | 0.496 | 6 | 0.307 | 2 | 0.679 | 6 |

The constancy of the candidate gene was ranked by the RefFinder program according to the integrating results of the ΔCt method, GeNorm, NormFinder, and BestKeeper. According to RefFinder, the stability of genes among the developmental stages was ranked as follows: *SOD* > *GAPDH* > *ACTIN* > *RPL10a* > *FOXO* > *RPL28* > *RPL18* > *RPS15* > *RPS13* (Fig. 2a). The GeNorm analysis suggested that all values of Vn/Vn+1 were above 0.15. Therefore, three reference genes should be used when normalizing the expression of hypothetical target gene in *T. dendrolimi* during qPCR analysis. The top three stable genes, *SOD, GAPDH*, and *ACTIN*, formed the most appropriate combination of reference genes for the adjustment of errors among developmental stages (Table 3).

**Fig. 2.**
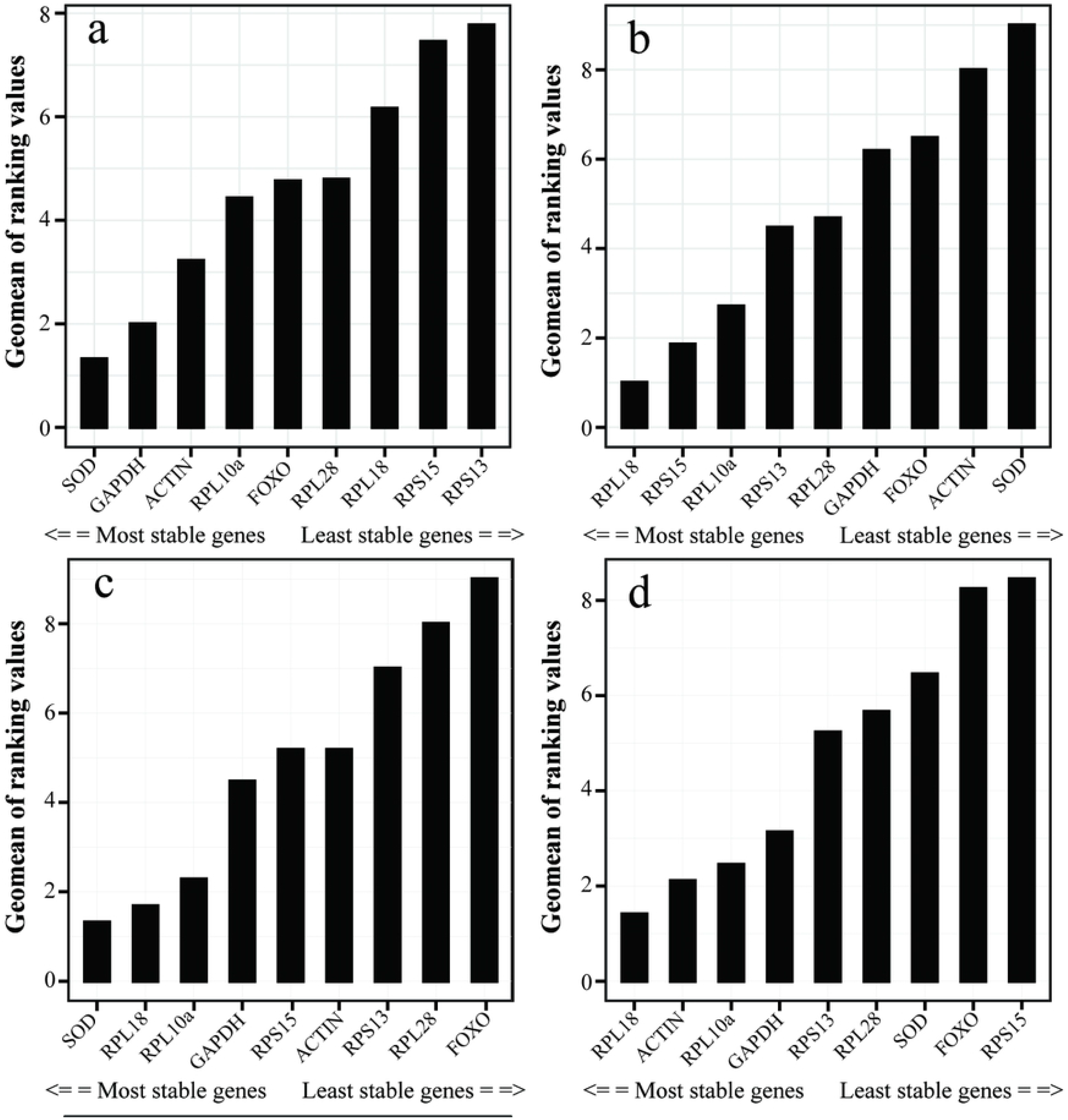
The constancy of the expression levels of nine candidate reference genes of *T. dendrolimi* across different developmental stage (a), tissue (b), sexes of adult (c), and temperature (d) based on RefFinder stability value.

**Table 3.** Recommended reference genes for different factor.

| Experimental Conditions | Reference Genes |  |  |
| --- | --- | --- | --- |
| **Developmental stage** | *SOD* | *GAPDH* | *ACTIN* |
| **Tissue** | *RPL18* | *RPS15* | — |
| **Sex** | *SOD* | *RPL18* | — |
| **Temperature** | *RPL18* | *RPL10* | — |

#### Tissue

According to the four algorithms, the top three stable genes affected by differences in tissue were *RPL18, RPS15*, and *RPL10a*, and the top two unstable genes were *SOD* and *ACTIN*. The third lowest stable gene was *FOXO* according to GeNorm and BestKeeper, or *GAPDH* according to the ΔCt method and NormFinder (Table 2).

According to RefFinder, the stability of expression among tissues was ranked as follows by gene: *RPL18* > *RPS15* > *RPL10a* > *RPS13* > *RPL28* > *GAPDH* > *FOXO* > *ACTIN* > *SOD* (Fig. 2b). According to results of GeNorm, the values of V2/3 were below 0.15. Therefore, the combination of two reference genes is recommended for adjusting the expression of genes in different tissues of *T. dendrolimi* (Fig. 3). The top two stable genes, *RPL18* and *RPS15*, formed the most appropriate combination of reference genes for the adjustment of errors among tissues (Table 3).

**Fig. 3.**
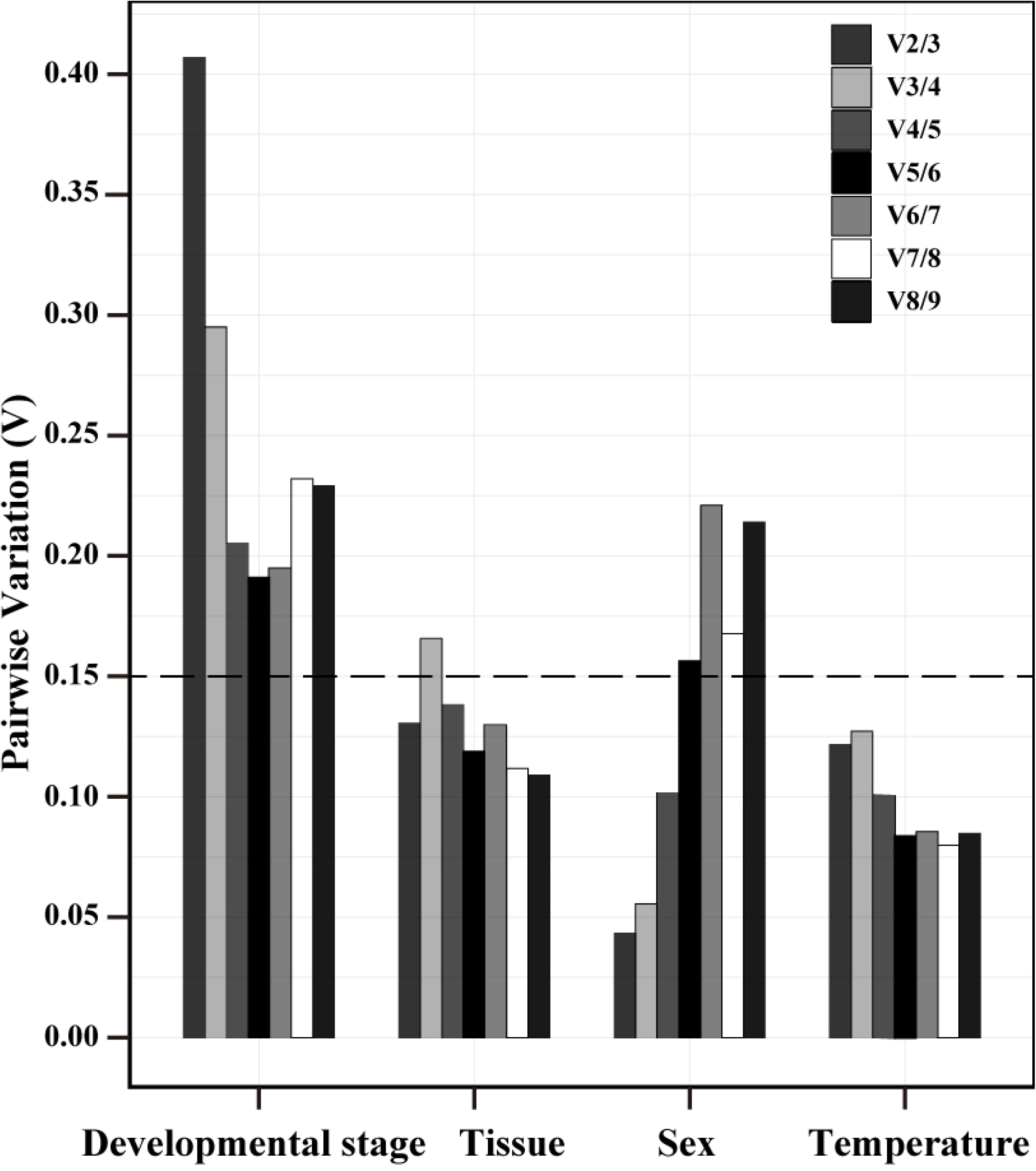
Pairwise variation (Vn/Vn+1) analysis of the candidate reference genes as influenced by different factors.

#### Sex

According to the four algorithms, the top three stable genes between females and males were *SOD, RPL18*, and *RPL10a*, and the top three unstable genes were *FOXO, RPL28*, and *RPS13* (Table 2).

According to RefFinder, the stability of expression as influenced by sex were ranked as follows by gene: *SOD* > *RPL18* > *RPL10a* > *GAPDH* > *ACTIN* = *RPS15* > *RPS13* > *RPL28* > *FOXO* (Fig. 2c). The analysis of GeNorm showed the values of V2/3 were below 0.15. Therefore, the combination of at least two reference genes is recommended for normalizing the expression of hypothetical target genes as influenced by sex (Fig. 3). *SOD* and *RPL18* formed the most suitable combination of reference genes for the adjustment of errors induced by sex (Table 3).

#### Temperature

According to the ΔCt method, GeNorm, and NormFinder, the top three genes in terms of expression stability under varying temperature were *RPL18, RPL10a*, and *RPS13*, and the top three unstable genes were *ACTIN, FOXO*, and *RPS15*. According to BestKeeper, the top three stable genes were *RPL18, SOD*, and *GAPDH*, and the top three unstable genes were *RPL28, FOXO*, and *RPS15* (Table 2).

According to RelFinder, the stability of expression as influenced by temperature were ranked as follows by gene: *RPL18* > *RPL10a* > *RPS13* > *GAPDH* > *SOD* > *RPL28* > *RPS15* > *FOXO* > *ACTIN* (Fig. 2d). The results of GeNorm showed that the values of V2/3 were below 0.15. Therefore, the combination of two reference genes should be used for normalizing the expression of target genes as influenced by temperature (Fig. 3). *RPL18* and *RPL10a* formed the most suitable combination of reference genes for the adjustment of errors induced by temperature (Table 3).

### Validation of the selected reference genes

To validate the accuracy of the recommended reference gene under different temperature conditions, *HSP90* was selected as a hypothetical target gene. The expression of *HSP90* was adjusted by applying the most stable gene, *RPL18*; the most suitable combination of the top two stable genes, *RPL18* and *RPL10a*; or the most unstable gene, *ACTIN*, and the second most unstable gene, *FOXO*, for normalization. Regardless of the different reference genes used, the relative expression level of *HSP90* at 32 °C was significantly higher than that at 25 °C. When the results were adjusted by using *RPL18* or the combination of *RPL18* and *RPL10a*, the relative expression level of *HSP90* at 32°C or 17°C was significantly higher than that at 25°C. However, when the value was normalized by *ACTIN* or *FOXO*, the results were not similar to that normalized by the recommended reference gene, RPL18, or the combination of *RPL18* and *RPL10a*. When *HSP90* expression was normalized by *ACTIN*, the difference between the results for 17°C and 32°C was insignificant. There was also no significant difference between 17 °C and 25 °C for results of *HSP90* expression normalized by *FOXO* (Fig. 4).

**Fig. 4.**
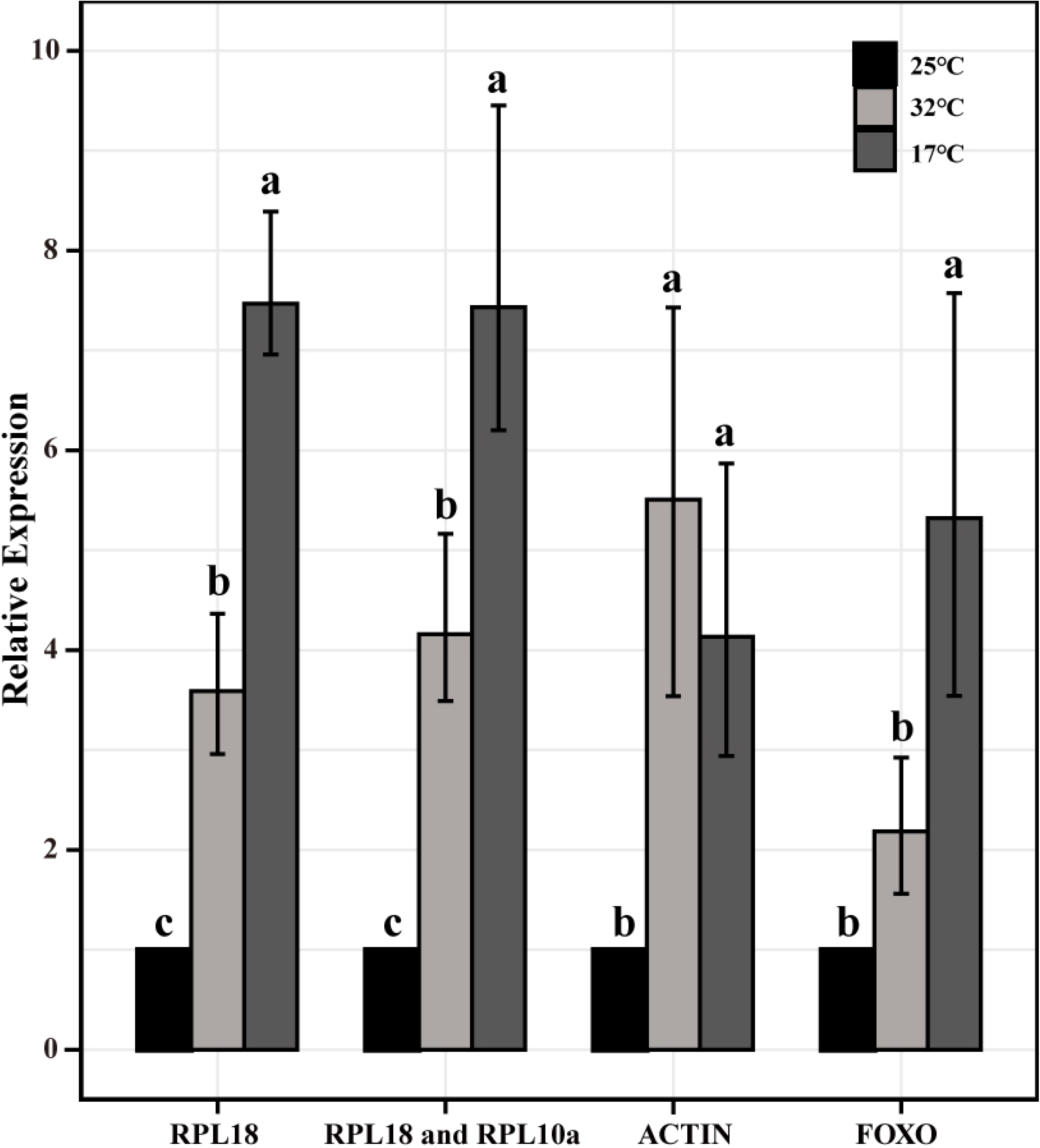
Relative expression levels of *HSP90* (MEAN ± SEM) under temperature in *T. dendrolimi*. normalized by stable reference genes (*RPL18* and the combination of *RPL18* and *RPL10*), or unstable genes (*ACTIN* and *FOXO*). Different letters indicate significant difference between the value normalized by different genes according to Tukey-Kramer test (*P* < 0.05).

## Discussion

The qPCR analysis is viewed as the model methodology to measure the expression of genes [46,47]. The selection of suitable reference genes is particularly important for qPCR procedure [33,34]. Thus far, the selection of reference genes for robust qPCR haven’t been reported at all for the *Trichogramma* genus. As we outlined in the introduction, the quality and quantity of the RNA sample of these tiny parasitoids are easily affected by biotic or abiotic factors. In present study, when the value of the hypothetical target gene, *HSP90*, was normalized using unsuitable genes, the results were different to those adjusted using the recommended reference genes (Fig. 4). This implies that the application of inappropriate reference genes reduces the repeatability of experiments and lead to misleading conclusions.

In present study, four different algorithms, the ΔCt method, NormFinder, GeNorm, and BestKeeper, were used to identify the stability of nine candidate genes. Although these algorithms produced globally similar rankings, some differences could be observed due to the different criteria of algorithms. These discrepancies arise from the specific working hypotheses and different premises of the algorithm. However, the same general pattern could be observed across the different algorithms under most conditions.

In our study, the *RPL18* gene was selected as a tissue-specific, temperature-specific, and sex-specific reference gene for normalization in the qPCR procedure. *RPL10a* and *RPS15* were the second most stable genes at different temperatures and tissues, respectively. The gene family of ribosomal proteins is the component of ribosomes and is constantly expressed throughout all developmental stages and in all living cells of the organisms [25]. Some of the genes have been used for normalizing the expression level of target genes in Hymenopteran insects and other insects, such as *A. mellifera* (*RP49, RPL13a*, and *RPS18*) [34], *L. japonica* (*RPL18, RPL27*, and *RPS18*) [20], *Solenopsis invicta* (*RPL18*) [48], *Tetranychus cinnabarinus* (*RPL13a*, and *RPS18*) [22], and *Holotrichia oblita* (*RPL13a, RPL18, RPS18, RPS3*, and *RPS6*) [49].

Regarding the scavenging of reactive oxygen species in organisms, *SOD* is considered a key gene expressing a product involved in this process [50]. In our study, *SOD* was identified as a developmental stage-specific and sex-specific reference gene, but it was not stable genes across different tissue samples and temperature conditions. In previous studies, the constancy of *SOD* was variable under different treatments in the organisms. For example, *SOD* was one of the most stable genes for different sexes of adults and developmental stages of *Spodoptera exigua*, whereas its expression was unstable across different tissues and specific larval stages [26]. The expression of *SOD* was stable across different photoperiod in *Harmonia axyridis*; however, its expression was unstable across different different tissues, different sexes of adults, developmental stages, and different temperature [51].

Aside from *SOD*, the candidate genes, *GAPDH* and *ACTIN*, were the 2nd and 3rd most stable genes across different developmental stages, respectively. *GAPDH*, which is important for energy metabolism in organisms, is a popular reference gene applied in qPCR studies of *A. mellifera* [34], *S. invicta* [47], *B. terrestris* [32], and *B. lucorum* [32]. *ACTIN* encodes proteins that are major contributors to the contractile property of muscles and cells and is expressed throughout all developmental stages and tissues of organisms [39]. It has been considered the ideal candidate reference gene for qPCR procedure and has been investigated extensively [25]. Many studies have shown that *ACTIN* expressed stable under the different developmental stages of many insects [25], including *Schistocerca gregaria* [52], *A. mellifera* [34], *Drosophila melanogaster* [53], *Chilo suppressalis* [54], *Chortoicetes terminifera* [55], *Liriomyza trifolii* [56], and *Diuraphis noxia* [57]. Nevertheless, the expression of *ACTIN* was not stable in several species of insects, including *Bemisia tabaci* [39], *Nilaparvata lugens* [27], and *Dendroctonus valens* [58].

## Conclusions

We evaluated nine candidate reference genes in *T. dendrolimi* under different conditions. The appropriate combination of reference genes should involve at least three genes for different developmental stages, two genes for different tissues, two genes for different sexes, and two genes for different temperatures, respectively. We identified the developmental stage-specific reference genes, *SOD, GAPDH*, and *ACTIN*; tissue-specific reference genes, *RPL18* and *RPS15*; sex-specific reference genes, *SOD* and *RPL18*; and temperature-specific reference genes, *RPL18* and *RPL10*. The application of the stable vs. unstable reference genes cause substantial differences in the expression level of a hypothetical target gene, *HSP90*. The results provide a basic reference for the robust procedure of qPCR for the quantification of gene expression in *T. dendrolimi*.

## Acknowledgments

We should thank ELSEVIER Language Editing Services for improving the language. This research was funded by the Projects of Guizhou Tobacco Corporation (201936, 201937, and 201941), the Major Projects of China National Tobacco Corporation (110202001032 (LS-01)), National Natural Science Foundation of China (31972339), the Agricultural Science and Technology Innovation Program (CAAS-ZDRW202108), Natural Science Foundation of Liaoning Province (2020-BS-137), and the Fundamental Research Funds for the Universities of Liaoning Province (LR2019061)

## Author Contributions

**Conceptualization:** Hui Dong, Li-sheng Zhang, Wu-nan Che.

**Data curation:** Liang-xiao Huo, Xue-ping Bai.

**Formal analysis:** Liang-xiao Huo, Jin-cheng Zhou.

**Funding acquisition:** Hui Dong.

**Investigation:** Xue-ping Bai, Su -fang Ning, Lin Lv.

**Writing – original draft:** Liang-xiao Huo, Jin-cheng Zhou.

**Writing – review & editing:** Jin-cheng Zhou, Li-sheng Zhang, Hui Dong.

## References

1. Zhou JC, Dong QJ, Zhang TS, Duan LJ, Ning SF, Liu QQ, et al. Effect of wind time on the dispersal capacity of Trichogramma dendrolimi Matsumura (Hymenoptera Trichogrammatidae). J. Asia. Pac. Entomol. 2019; 22: 742–749. https://doi.org/10.1016/j.aspen.2019.06.001

2. Zang LS, Wang S, Zhang F, Desneux N. Biological Control with Trichogramma in China: History, Present Status, and Perspectives. Annu. Rev. Entomol. 2021; 66: 463–484. https://doi.org/10.1146/annurev-ento-060120-091620 PMID: 32976724

3. Smith SM. Biological control with Trichogramma: advances, successes, and potential of their use. Annu. Rev. Entomol. 1996; 41: 375–406. https://doi.org/10.1146/annurev.en.41.010196.002111 PMID: 15012334

4. Luo S, Naranjo SE, Wu K. Biological control of cotton pests in China. Biol. Control 2014; 68: 6–14. http://dx.doi.org/10.1016/j.biocontrol.2013.06.004

5. Huang J, Zhang B, Zhang F, Li YX. Effect of rearing host on the size and egg load of three Trichogramma species (Hymenoptera: Trichogrammatidae). Acta Entomol. Sin. 2015; 58: 1098–1107. http://dx.doi.org/10.16380/j.kcxb.2015.10.008

6. Liu QQ, Zhou JC, Zhang C, Ning SF, Duan LJ, Dong H. Co-occurrence of thelytokous and bisexual Trichogramma dendrolimi Matsumura (Hymenoptera: Trichogrammatidae) in a natural population. Sci. Rep. 2019; 9: 17480. https://doi.org/10.1038/s41598-019-53992-8 PMID: 31767914

7. Zhang J, Ruan C, Zang L, Shao X, Shi S. Technological improvements for mass production of Trichogramma and current status of their applications for biological control on agricultural pests in China. Chinese J. Biol. Control 2015; 31: 638–646. https://doi.org/10.16409/j.cnki.2095-039x.2015.05.004

8. Zhang JJ, Zhang X, Zang LS, D. WM, Hou YY, Ruan CC, et al. Advantages of diapause in Trichogramma dendrolimi mass production via eggs of the Chinese silkworm, Antheraea pernyi. Pest Manag. Sci. 2018; 74: 959–965. https://doi.org/10.1002/ps.4795 PMID: 29155485

9. Li TH, Tian CY, Zang LS, Hou YY, Ruan CC Yang X, et al. Multiparasitism with Trichogramma dendrolimi on egg of Chinese oak silkworm, Antheraea pernyi, enhances emergence of Trichogramma ostriniae. J. Pest Sci. 2019; 92: 707–713. https://doi.org/10.1007/s10340-018-1018-5

10. Zhou JC, Liu QQ, Wang QR, Ning SF, Che WN, Dong H. Optimal clutch size for quality control of bisexual and Wolbachia-infected thelytokous lines of Trichogramma dendrolimi Matsumura (Hymenoptera: Trichogrammatidae) mass reared on eggs of a substitutive host, Antheraea pernyi Guérin-Méneville (Lepidoptera: Saturniidae). Pest Manag. Sci. 2020; 76: 2635–2644. https://doi.org/10.1002/ps.5805 PMID: 32112519

11. Tian JC, Wang ZC, Wang GR, Zhong LQ, Zheng XS, Xu HX, et al. The Effects of Temperature and Host Age on the Fecundity of Four Trichogramma Species, Egg Parasitoids of the Cnaphalocrocis medinalis (Lepidoptera: Pyralidae). J. Econ. Entomol. 2017; 110: 949–953. https://doi.org/10.1093/jee/tox108 PMID:28398560

12. Lindsey A, Kelkar YD, Wu X, Sun D, Martinson EO, Yan Z, et al. Comparative genomics of the miniature wasp and pest control agent Trichogramma pretiosum. BMC Biol. 2018; 16: 54. https://doi.org/10.1186/s12915-018-0520-9 PMID: 29776407

13. Ferguson KB, Kursch-Metz T, Verhulst EC, Pannebakker BA. Hybrid genome assembly and evidence-based annotation of the egg parasitoid and biological control agent Trichogramma brassicae. G3 (Bethesda) 2020; 10: 3533–3540. https://doi.org/10.1534/g3.120.401344 PMID: 32792343

14. Ferguson BK. Trichogramma evanescens, whole genome shotgun sequencing project. 2020; Available from: https://www.ncbi.nlm.nih.gov/genome/87258.

15. VanGuilder HD, Vrana KE, Freeman WM. Twenty-five years of quantitative PCR for gene expression analysis. Biotechniques 2008; 44: 619–626. https://doi.org/10.2144/000112776 PMID: 18474036

16. Yan JW, Yuan FR, Long GY, Qin L, Deng ZN. Selection of reference genes for quantitative real-time RT-PCR analysis in citrus. Mol Biol Rep. 2012; 39: 1831– 1838. https://doi.org/10.1007/s11033-011-0925-9 PMID: 21633888

17. Andersen CL, Jensen JL, Ørntof TF. Normalization of real-time quantitative reverse transcription-PCR data: a model-based variance estimation approach to identify genes suited for normalization, applied to bladder and colon cancer data sets. Cancer Res. 2004; 64: 5245–5250. https://doi.org/10.1158/0008-5472.CAN-04-0496 PMID: 15289330

18. Gao ZH, Deng WH, Zhu F. Reference gene selection for quantitative gene expression analysis in black soldier fly (Hermetia illucens). PLoS ONE 2019; 14: e0221420. https://doi.org/10.1371/journal.pone.0221420 PMID: 31419256

19. Gao XK, Zhang S, Luo JY, Wang CY, Lü Lm, Zhang LJ, et al. Comprehensive evaluation of candidate reference genes for gene expression studies in Lysiphlebia japonica (Hymenoptera: Aphidiidae) using RT-qPCR. Gene 2017a; 637: 211–218. https://doi.org/10.1016/j.gene.2017.09.057 PMID: 28964897

20. Gao XK, Zhang S, Luo JY, Wang CY, Lü LM, Zhang LJ, et al. Identification and validation of reference genes for gene expression analysis in Aphidius gifuensis (Hymenoptera: Aphidiidae). PLoS ONE 2017b; 12: e0188477. https://doi.org/10.1371/journal.pone.0188477 PMID: 29190301

21. Jian B, Liu B, Bi YR, Hou WS, Wu CX, Han TF. Validation of internal control for gene expression study in soybean by quantitative real-time PCR. BMC Mol. Biol. 2008; 9: 1–14. https://doi.org/10.1186/1471-2199-9-59 PMID: 18573215

22. Sun W, Jin Y, He L, Lu WC, Li M. Suitable reference gene selection for different strains and developmental stages of the carmine spider mite, Tetranychus cinnabarinus, using quantitative real-time PCR. J. Insect Sci. 2010; 10: 208. https://doi.org/10.1673/031.010.20801 PMID: 21265619

23. Dzaki N, Ramli KN, Azlan A, Ishak IH, Azzam G. Evaluation of reference genes at different developmental stages for quantitative real-time PCR in Aedes aegypti. Sci. Rep. 2017; 7: 43618. https://doi.org/10.1038/srep43618 PMID: 28300076

24. Chapman JR, Waldenström, J. With Reference to Reference Genes: A Systematic Review of Endogenous Controls in Gene Expression Studies. PLoS ONE 2015; 10: e0141853. https://doi.org/10.1371/journal.pone.0141853 PMID: 26555275

25. Lü J, Yang C, Zhang Y, Pan H. Selection of Reference Genes for the Normalization of RT-qPCR Data in Gene Expression Studies in Insects: A Systematic Review. Front. Physiol. 2018; 9: 1560. https://doi.org/10.3389/fphys.2018.01560 PMID: 30459641

26. Zhu X, Yuan M, Shakeel M, Zhang Y, Wang S, Wang X, et al. Selection and evaluation of reference genes for expression analysis using qRT-PCR in the beet armyworm Spodoptera exigua (Hübner) (Lepidoptera: Noctuidae). PLoS ONE 2014; 9: e84730. https://doi.org/10.1371/journal.pone.0084730 PMID: 24454743

27. Yuan M, Lu Y, Zhu X, Wan H, Shakeel M, Zhan S, et al. Selection and evaluation of potential reference genes for gene expression analysis in the brown planthopper, Nilaparvata lugens (Hemiptera: Delphacidae) using reverse-transcription quantitative PCR. PLoS ONE 2014; 9: e86503. https://doi.org/10.1371/journal.pone.0086503 PMID: 24466124

28. Ponton F, Chapuis MP, Pernice M, Sword GA, Simpson SJ. Evaluation of potential reference genes for reverse transcription-qPCR studies of physiological responses in Drosophila melanogaster. J. Insect Physiol. 2011; 57: 840–850. https://doi.org/10.1016/j.jinsphys.2011.03.014 PMID: 21435341

29. Dong M, Zhang X, Chi X, Mou S, Xu J, Xu D, et al. The validity of a reference gene is highly dependent on the experimental conditions in green alga Ulva linza. Curr. Genet. 2012; 58: 13–20. https://doi.org/10.1007/s00294-011-0361-3 PMID: 22205301

30. Li QY, Li ZL, Lu MX, Cao SS, D. YZ. Selection of valid reference genes for quantitative real-time PCR in Cotesia chilonis (Hymenoptera: Braconidae) exposed to different temperatures. PLoS ONE 2019; 14: e0226139. https://doi.org/10.1371/journal.pone.0226139 PMID: 31877150

31. Niu J, Cappelle K, de Miranda JR, Smagghe G, Meeus, I. Analysis of reference gene stability after Israeli acute paralysis virus infection in bumblebees Bombus terrestris. J. Invertebr. Pathol. 2014, 115, 76–79. https://doi.org/10.1016/j.jip.2013.10.011 PMID: 24184950

32. Hornáková D, Matousková P, Kindl J, Valterová I, Pichová I. Selection of reference genes for real-time polymerase chain reaction analysis in tissues from Bombus terrestris and Bombus lucorum of different ages. Anal. Biochem. 2010; 397: 118–120. https://doi.org/10.1016/j.ab.2009.09.019 PMID: 19751695

33. Freitas F, Depintor TS, Agostini LT, Luna-Lucena D, Nunes F, Bitondi M, et al. Evaluation of reference genes for gene expression analysis by real-time quantitative PCR (qPCR) in three stingless bee species (Hymenoptera: Apidae: Meliponini). Sci. Rep. 2019; 9: 17692. https://doi.org/10.1038/s41598-019 53544-0 PMID: 31776359

34. Scharlaken B, de Graaf DC, Goossens K, Brunain M, Peelman LJ, Jacobs FJ. Reference gene selection for insect expression studies using quantitative real-time PCR: The head of the honeybee, Apis mellifera, after a bacterial challenge. J. Insect Sci. 2008; 8: 1–10. http://dx.doi.org/10.1673/031.008.3301

35. Xu J, Welker DL, James RR. Variation in Expression of Reference Genes across Life Stages of a Bee, Megachile rotundata. Insects 2021; 12: 36. https://doi.org/10.3390/insects12010036 PMID: 33418888

36. Wang XJ. The effect of temperature and host on the body size of Trichogramma dendrolimi (Matsumura) and expression level of FoxO gene. Master Thesis, Shenyang Agricultural University. 2016. Available from: https://kns.cnki.net/kcms/detail/detail.aspx?FileName=1016146291.nh&DbName=CMFD2017

37. Liu QQ, Zhang C, Zhou JC, Dong QJ, Huo LX, Dong H. One simple, rapid and economical method for ploidy detection of Trichogramma dendrolimi Matsumura. J. Asia-Pac Entomol. 2020; 23: 345–349. https://doi.org/10.1016/j.aspen.2019.12.010

38. Zhang L, Huang J, Dong X, Zhang F, Li Y. Superparasitism and Ontogeny of Two Trichogramma Species on Corcyra cephalonica (Stainton). Chinese J. Biol. Control 2015; 31: 481–486. https://doi.org/10.16409/j.cnki.2095-039x.2015.04.006

39. Li R, Xie W, Wang S, Wu Q, Yang N, Yang X, et al. Reference gene selection for qRT-PCR analysis in the sweetpotato whitefly, Bemisia tabaci (Hemiptera: Aleyrodidae). PLoS ONE 2013, 8, e53006. https://doi.org/10.1371/journal.pone.0053006 PMID: 23308130

40. Vandesompele J, Preter KD, Pattyn F, Poppe B, Roy NV, Paepe AD, et al. Accurate normalization of real-time quantitative RT-PCR data by geometric averaging of multiple internal control genes. Genome Biol. 2002; 3: RESEARCH0034. https://doi.org/10.1186/gb-2002-3-7-research0034 PMID: 12184808

41. Pfaffl MW, Tichopad A, Prgomet C, Neuvians TP. Determination of stable housekeeping genes, differentially regulated target genes and sample integrity: BestKeeper--Excel-based tool using pair-wise correlations. Biotechnol. Lett. 2004; 26: 509–515. https://doi.org/10.1023/b:bile.0000019559.84305.47 PMID: 15127793

42. Silver N, Best S, Jiang J, Thein SL. Selection of housekeeping genes for gene expression studies in human reticulocytes using real-time PCR. BMC Mol. Biol. 2006; 7: 33. https://doi.org/10.1186/1471-2199-7-33 PMID: 17026756

43. Xie F, Xiao P, Chen D, Xu L, Zhang, B. miRDeepFinder: a miRNA analysis tool for deep sequencing of plant small RNAs. Plant Mol. Biol. 2012; 80: 75–84. https://doi.org/10.1007/s11103-012-9885-2 PMID: 22290409

44. Livak KJ, Schmittgen TD. Analysis of relative gene expression data using real-time quantitative PCR and the 2(-Delta Delta C(T)) Method. Methods 2001; 25: 402–408. https://doi.org/10.1006/meth.2001.1262 PMID: 11846609

45. R Core Team. 2021. R: A language and environment for statistical computing. R Foundation for Statistical Computing, Vienna, Austria.2021. URL https://www.R-project.org/

46. Ponchel F, Toomes C, Bransfield K, Leong FT, Douglas SH, Field SL, et al. Real-time PCR based on SYBR-green I fluorescence: An alternative to the taqman assay for a relative quantification of gene rearrangements, gene amplifications and micro gene deletions. BMC Biotechnol. 2003; 3: 18. https://doi.org/10.1186/1472-6750-3-18 PMID: 14552656

47. Bao W, Qu Y, Shan X, Wan Y. Screening and Validation of Housekeeping Genes of the Root and Cotyledon of Cunninghamia lanceolata under Abiotic Stresses by Using Quantitative Real-Time PCR. Int. J. Mol. Sci. 2016; 17: 1198. https://doi.org/10.3390/ijms17081198 PMID: 27483238

48. Cheng D, Zhang Z, He X, Liang G. Validation of reference genes in Solenopsis invicta in different developmental stages, castes and tissues. PLoS ONE 2013; 8: e57718. https://doi.org/10.1371/journal.pone.0057718 PMID: 23469057

49. Xie M, Zhong Y, Lin L, Zhang G, Su W, Ni W, et al. Evaluation of reference genes for quantitative real-time PCR normalization in the scarab beetle Holotrichia oblita. PLoS ONE 2020; 15: e0240972. https://doi.org/10.1371/journal.pone.0240972 PMID: 33085726

50. Xikeranmu Z, Abdunasir M, Ma J, Tusong K, Liu XN. Characterization of two copper/zinc superoxide dismutases (Cu/Zn-SODs) from the desert beetle Microdera punctipennis and their activities in protecting E. coli cells against cold. Cryobiology 2019; 87: 15–27. https://doi.org/10.1016/j.cryobiol.2019.03.006 PMID: 30890324

51. Qu C, Wang R, Che W, Zhu X, Li F, Luo C. Selection and evaluation of reference genes for expression analysis using quantitative real-time PCR in the Asian Ladybird Harmonia axyridis (Coleoptera: Coccinellidae). PLoS ONE 2018; 13: e0192521. https://doi.org/10.1371/journal.pone.0192521 PMID: 29889877

52. Van Hiel MB, Van Wielendaele P, Temmerman L, Van Soest S, Vuerinckx K, Huybrechts R, et al. Identification and validation of housekeeping genes in brains of the desert locust Schistocerca gregaria under different developmental conditions. BMC Mol. Biol. 2009; 10: 56. https://doi.org/10.1186/1471-219910-56 PMID: 19508726

53. Ponton F, Chapuis MP, Pernice M, Sword GA, Simpson SJ. Evaluation of potential reference genes for reverse transcription-qPCR studies of physiological responses in Drosophila melanogaster. J. Insect Physiol. 2011; 57: 840–850. https://doi.org/10.1016/j.jinsphys.2011.03.014 PMID: 21435341

54. Teng X, Zhang Z, He G, Yang L, Li F. Validation of reference genes for quantitative expression analysis by real-time RT-PCR in four Lepidopteran insects. J. Insect Sci. 2012; 12: 60. https://doi.org/10.1673/031.012.6001 PMID: 22938136

55. Chapuis MP, Tohidi-Esfahani D, Dodgson T, Blondin L, Ponton F, Cullen D, et al. Assessment and validation of a suite of reverse transcription-quantitative PCR reference genes for analyses of density-dependent behavioural plasticity in the Australian plague locust. BMC Mol. Biol. 2011; 12: 7. https://doi.org/10.1186/1471-2199-12-7 PMID: 21324174

56. Chang YW, Chen JY, Lu MX, Gao Y, Tian ZH, Gong WR, et al. Selection and validation of reference genes for quantitative real-time PCR analysis under different experimental conditions in the leafminer Liriomyza trifolii (Diptera: Agromyzidae). PLoS ONE 2017; 12: e0181862. https://doi.org/10.1371/journal.pone.0181862 PMID: 28746411

57. Sinha DK, Smith CM. Selection of reference genes for expression analysis in Diuraphis noxia (Hemiptera: Aphididae) fed on resistant and susceptible wheat plants. Sci. Rep. 2014; 4: 5059. https://doi.org/10.1038/srep05059 PMID: 24862828

58. Zheng C, Zhao D, Xu Y, Shi F, Zong S, Tao J. Reference Gene Selection for Expression Analyses by qRT-PCR in Dendroctonus valens. Insects 2020; 11: 328. https://doi.org/10.3390/insects11060328 PMID: 32471281

